# Predicting Knee Adduction Moment Response to Gait Retraining with Minimal Clinical Data

**DOI:** 10.1101/2021.09.29.462292

**Authors:** Nataliya Rokhmanova, Katherine J. Kuchenbecker, Peter B. Shull, Reed Ferber, Eni Halilaj

**Affiliations:** Mechanical Engineering, Carnegie Mellon University, Pittsburgh, PA, USA; Max Planck Institute for Intelligent Systems, Stuttgart, Germany; Mechanical Engineering, Shanghai Jiao Tong University, Shanghai, China; University of Calgary, Calgary, Alberta, Canada; Mechanical Engineering, Biomedical Engineering, the Robotics Institute, Carnegie Mellon University, Pittsburgh, PA, USA

## Abstract

Knee osteoarthritis is a progressive disease mediated by high joint loads. Foot progression angle modifications that reduce the knee adduction moment (KAM), a surrogate of knee loading, have demonstrated efficacy in alleviating pain and improving function. Although changes to the foot progression angle are overall beneficial, KAM reductions are not consistent across patients. Moreover, customized interventions are time-consuming and require instrumentation not commonly available in the clinic. We present a model that uses minimal clinical data to predict the extent of first peak KAM reduction after toe-in gait retraining. For such a model to generalize, the training data must be large and variable. Given the lack of large public datasets that contain different gaits for the same patient, we generated this dataset synthetically. Insights learned from ground-truth datasets with both baseline and toe-in gait trials (N=12) enabled the creation of a large (N=138) synthetic dataset for training the predictive model. On a test set of data collected by a separate research group (N=15), the first peak KAM reduction was predicted with a mean absolute error of 0.134% body weight * height (%BW*HT). This error is smaller than the test set’s subject average standard deviation of the first peak during baseline walking (0.306 %BW*HT). This work demonstrates the feasibility of training predictive models with synthetic data and may provide clinicians with a streamlined pathway to identify a patient-specific gait retraining outcome without requiring gait lab instrumentation.

**Author Summary:** Gait retraining as a conservative intervention for knee osteoarthritis shows great promise in extending pain-free mobility and preserving joint health. Although customizing a treatment plan for each patient may help to ensure a therapeutic response, this procedure cannot yet be performed outside of the gait laboratory, preventing research advances from becoming a part of clinical practice. Our work aims to predict the extent to which a patient with knee osteoarthritis will benefit from a non-invasive gait retraining therapy using measures that can be easily collected in the clinic. To overcome a lack of normative databases for gait retraining, we generated data synthetically based on limited ground-truth examples, and provided experimental evidence for the model’s ability to generalize to new subjects by evaluating on data collected by a separate research group. Our results can contribute to a future in which predicting the therapeutic benefit of a potential treatment can determine a custom treatment path for any patient.

## 1. Introduction

Globally, one in five individuals aged 40 and older are afflicted by knee osteoarthritis, a painful joint disease that still lacks a cure or clinically accepted disease-modifying intervention [1]. Pain is managed pharmaceutically, while structurally cartilage is left to degrade until joint failure, at which point joint replacement surgery is recommended. Although the etiology of the disease is still an active area of interdisciplinary investigation, disease progression is known to be exacerbated by high joint loading during ambulation [2]. Since the forces that stress the tibiofemoral contact surface generally cannot be measured *in vivo*, the knee adduction moment (KAM) is often used as a surrogate of loading on the medial compartment of the knee [3]. Both a higher peak KAM [4] and higher KAM impulse [5] are associated with osteoarthritis progression: reducing either or both peaks of the typically two-peaked KAM has therefore been a primary target of non-invasive gait retraining interventions [6].

Biomechanists continue to direct effort to advance conservative interventions and preserve native joint health to the greatest extent possible [7]. Gait modification strategies to reduce KAM have included decreasing walking speed [8,9], increasing trunk sway [8,10-13], or changing the foot progression angle (FPA) [14-19]. For some, the requisite decrease in walking speed can be prohibitive to daily living [9], and increasing trunk sway has been reported to induce back pain and imbalance [12]. In contrast, changing the foot progression angle successfully reduces KAM with minimal discomfort [20] and is generally preferred by patients [16]. Walking with the toes pointed inward can reduce the first peak of KAM by lateralizing the center of pressure and medializing the knee joint center (Fig 1). After six weeks of retraining a toe-in gait, knee osteoarthritis patients reduced the first peak of KAM by 0.44 %BW*HT on average, reporting reduced knee pain and improved function at a one-month follow-up [18]. This reduction was comparable to high tibial osteotomy [21] but without the risks associated with the surgery [22].

**Fig 1.**
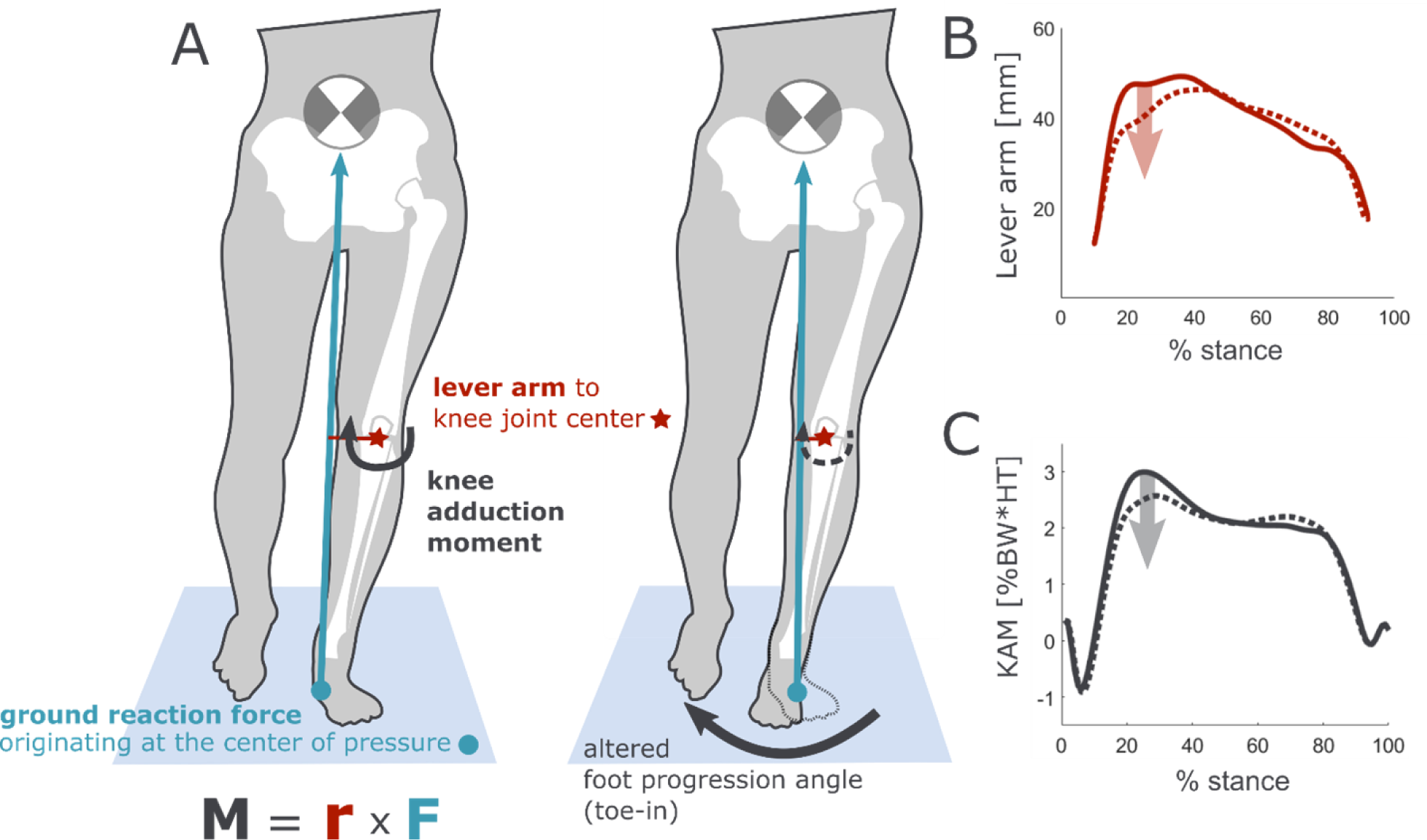
Toe-in gait reduces the knee adduction moment. (A) The moment about the knee is computed from the ground reaction force and lever arm to the knee joint center. Toe-in gait shifts the knee center medially and the foot center of pressure laterally in the first half of stance, reducing the (B) lever arm, which reduces (C) the KAM. Ground reaction force magnitude (not shown in the figure) does not change.

Customizing the change in foot progression angle to the individual results in a larger peak KAM reduction than a non-custom intervention [15]. However, determining a target foot progression angle is a time-consuming process of acclimation and evaluation that can be performed only in the gait lab. At present, researchers must iteratively evaluate KAM reduction at a range of foot progression angles; no methods currently exist to automate this procedure.

To extend gait lab advances into the clinic, we sought to build a predictive model for KAM reduction with toe-in gait using only features that can be easily obtained in the clinic (Fig 2). For such a model to be generalizable to new patients, large training data with both baseline and toe-in gait are needed, but this specialized dataset does not yet exist. To address this need, we synthesized toe-in gait from baseline walking data from 138 participants. The synthetic toe-in gait was generated using learned gait modification patterns extracted from a ground-truth dataset of 12 participants walking at baseline and toe-in. We evaluated the predictive model on an independent dataset collected in a different laboratory.

**Fig 2.**
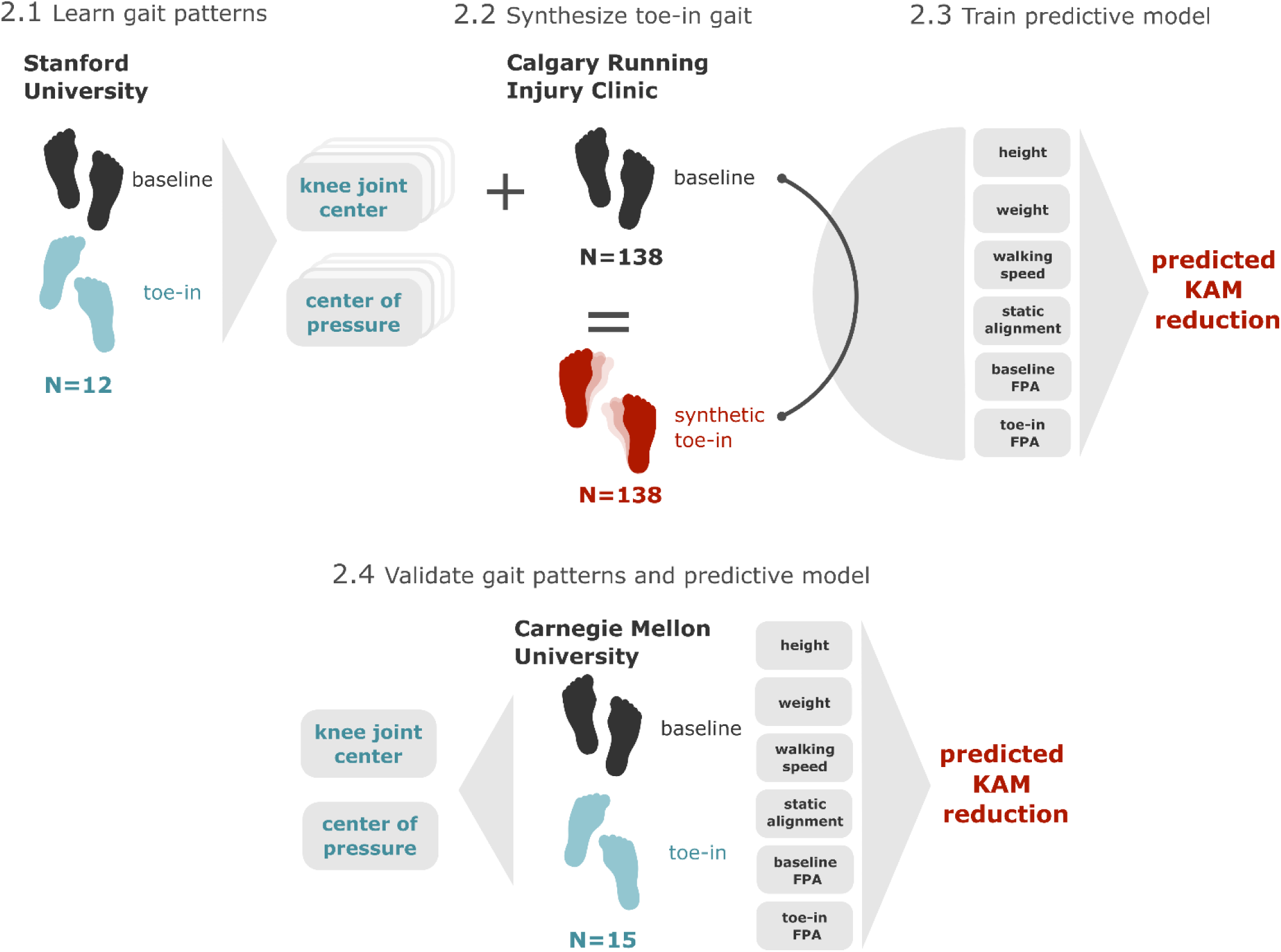
Predicting first peak KAM reduction. Two datasets are used to develop the toe-in gait data needed to train the predictive model: ground-truth data used to learn toe-in gait patterns across a range of toe-in angles and a large baseline database from which synthetic toe-in gait is generated. A set of six features predicts the reduction in the first peak of KAM. The predictive model, as well as the gait patterns of knee joint center and foot center of pressure locations with toe-in gait, are validated using an additional independent dataset.

## 2. Methods

### Ethics Statement

All experimental procedures involving human participants were approved by each institution’s ethics review boards: the Stanford University Institutional Review Board, the University of Calgary Conjoint Health Research Ethics Board, and the Carnegie Mellon University Institutional Review Board. All participants provided their written informed consent to participate.

#### 2.1 Learning gait patterns

Optical motion capture and force plate data were collected from 12 subjects with knee osteoarthritis (Table 1, Stanford University) and were used to learn how gait changes with increasing toe-in angle. Subjects walked at a self-selected speed on a split-belt instrumented treadmill under two conditions: walking normally (baseline gait) and walking with toes pointed in (toe-in gait). The last ten strides of each condition were used for analysis. We filtered force data at 15 Hz using a fourth order zero-lag Butterworth low-pass filter. Foot progression angle was defined in the laboratory horizontal plane as the angle between the anterior-posterior axis and the line connecting the markers placed on the calcaneus and the second metatarsal head. A toe-in angle was defined with respect to a participant’s average baseline foot progression angle. A specific toe-in angle was not enforced and mean and standard deviation (± STD) baseline and toe-in angles were 3.97° (± 4.91°) and -5.65° (± 4.10°) respectively.

**Table 1.** Summary demographics for participants included in each dataset

|  | **Stanford University** | **Calgary Running Injury Clinic** | **Carnegie Mellon University** |
| --- | --- | --- | --- |
| *Number of subjects* | 12 | 138 | 15 |
| *Sex* | 7 M / 5 F | 57 M / 81 F | 9 M / 6 F |
| *Height*<sup>1</sup> | 170.7 cm ($\pm 8.4$ cm) | 171.6 cm ($\pm 8.9$ cm) | 173.9 cm ($\pm 9.3$ cm) |
| *Weight*<sup>1</sup> | 77.7 kg ($\pm 18.0$ kg) | 70.2 kg ($\pm 12.4$ kg) | 68.0 kg ($\pm 11.1$ kg) |

<sup>1</sup> Mean ( ± STD)

First, at each step, the foot center of pressure and knee joint center in the mediolateral and anterior-posterior directions were expressed with respect to the pelvis center [16]. The pelvis center was calculated as the centroid of the markers placed on the left and right anterior and posterior superior iliac crests. We then normalized each step from heel strike to toe-off as 0 to 100% stance. For each subject, we represented the center of pressure and knee joint center trajectories during *toe-in gait* with respect to that subject’s average center of pressure and knee joint center trajectories during *baseline gait*. We labeled all 120 toe-in trajectories (10 steps x 12 subjects) by the toe-in angle at that step and grouped them into 1° bins from 1° to 10°. Within each bin, trajectories were averaged, creating a set of gait patterns across all subjects that we used to estimate the change in foot center of pressure and knee joint center for a given toe-in angle (Fig 3A). We represented each of the binned and averaged trajectories using spline bases of order 12 [23]. We then used these gait patterns to synthesize toe-in gait in new subjects by offsetting the learned center of pressure and knee joint center trajectory modifications for a given toe-in angle from the new subject’s baseline trajectories (Fig 3B).

**Fig 3.**
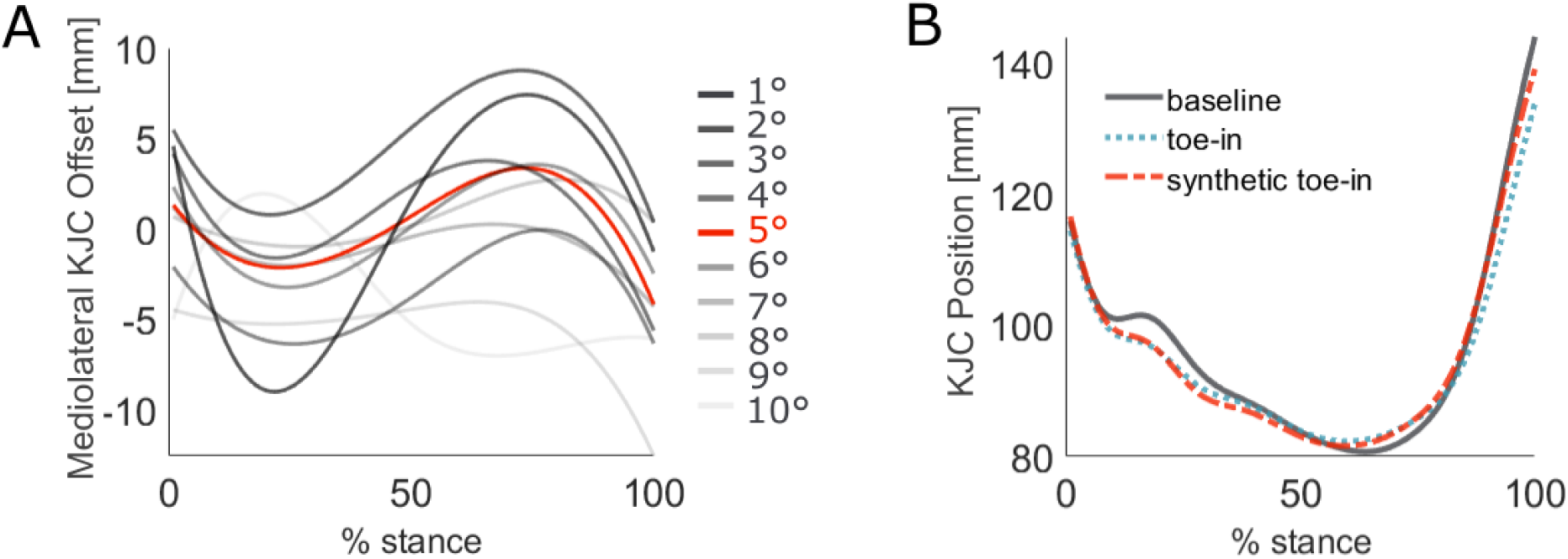
Synthetic gait generated via learned patterns. At each toe-in angle from 1° to 10°, all subject trajectories for the center of pressure and knee joint center (KJC) were binned and averaged. (A) At a given toe-in angle, the trajectory represents the positional offset from baseline gait. (B) Here, knee joint center position in the mediolateral direction was predicted for a representative subject with a 5° toe-in angle by adding the learned offset to the baseline trajectory.

To test whether the learned gait patterns were generalizable across subjects, we carried out exhaustive leave-one-out cross-validation by learning center of pressure and knee joint center trajectories from 11 subjects and testing on the left-out subject. We combined the synthesized center of pressure and knee joint center trajectories with the subject’s baseline ground reaction force to compute the KAM using the lever arm method [13]. We compared the synthetic KAM to the subject’s ground-truth toe-in KAM, computing the average root mean squared error (RMSE) (±STD) for the knee joint center, center of pressure, and KAM trajectories across all subjects. We assessed the accuracy of the first peak KAM estimate using the mean absolute error (MAE) (±STD) across all subjects.

#### 2.2 Synthesizing toe-in gait data

The learned gait patterns extracted from the ground-truth dataset were used to synthesize toe-in gait from an optical motion capture dataset of 138 subjects (Table 1, Calgary Running Injury Clinic). There were no exclusion criteria for participation, and some subjects were experiencing a lower extremity running-related injury, but all subjects were pain-free at the time of data collection. Subjects walked at a self-selected speed on an instrumented treadmill and data were collected for approximately 2 minutes. We filtered ground reaction force data at 15 Hz using a fourth-order zero-lag Butterworth low-pass filter and identified and removed steps that did not land cleanly on the force plates. We applied the toe-in gait patterns to the average and step-normalized knee joint center and center of pressure trajectories for the left leg of each subject, at each toe-in angle from 1° to 10°, resulting in a final dataset of 1380 entries (138 subjects x 10 toe-in angles). We computed the first peak of the KAM, scaled to percent body weight times height, for the baseline and synthetic toe-in gaits; the reduction in the first peak was used as the response variable of the predictive model.

#### 2.3 Training a predictive model

We used the synthetic toe-in gait data to train a predictive model for KAM reduction. To ensure that this model would be useful to clinicians without access to gait lab instrumentation, we used input features that are existing clinical measurements or can be computed with emerging wearable technologies. The features used were: height, weight, baseline walking speed, static knee alignment, baseline foot progression angle, and target toe-in angle. We computed static knee alignment as the angle between the vectors connecting 1) the lateral malleolus and lateral epicondyle and 2) the lateral epicondyle and hip joint center, in the frontal plane, with valgus angle defined as positive. These landmarks can be identified in the clinic by manual inspection and goniometric measurement and may be more accurately estimated in the future with video-based motion capture. Walking speed [24] and baseline foot progression angle [25,26] can be obtained with inertial measurement units, which are becoming more lightweight and commercially available.

We trained a linear regression model on the synthetic data to predict first peak KAM reduction. We split data by subject into 80%/10%/10% training, validation, and test sets, respectively. All input features were standardized to have a mean of zero and standard deviation of one using the training data. We used LASSO (Least Absolute Shrinkage and Selection Operator) [27] to identify the most relevant features that corresponded to the model with the smallest mean squared error (MSE). The alpha regularization term was tuned by training the model with five-fold cross-validation and selecting the alpha value that corresponded to the smallest MSE on the validation set. We selected alpha values from a range of 0 to 1 in increments of 0.1. To quantify model accuracy on synthetic data, we computed the MAE (±STD) between the synthetic first peak KAM reduction and the output of the predictive model. We also computed the mean signed error between the synthetic reduction and model prediction across all toe-in angles for each subject.

#### 2.4 Validating gait patterns and predictive model

To evaluate how well these methods would generalize to data collected in different settings, we tested the learned toe-in gait patterns and the resulting predictive model using newly collected gait data (Table 1, Carnegie Mellon University). Fifteen healthy participants walked at baseline and with toe-in gait on an instrumented treadmill at a self-selected speed for one minute during both conditions. Participants were guided to maintain a toe-in angle of approximately 5° relative to baseline with the use of vibration feedback on the shank [28], but average toe-in angles ranged from 3° to 10°. Mean and standard deviation (±STD) baseline and relative toe-in angles were 6.46° (±5.43°) and -6.60° (±1.72°), respectively. We computed synthetic toe-in center of pressure and knee joint center trajectories for each subject, using their average toe-in angle, their baseline gait, and the learned gait patterns from the Stanford ground-truth dataset, per the method described in Section 2.1.

The predictive model, trained on the synthetic data, was evaluated on the 15 new subjects using their height, weight, baseline walking speed, static knee alignment, baseline foot progression angle, and average toe-in angle as input features. We computed the MAE (±STD) between the real first peak KAM reduction and the output of the predictive model, as well as the mean signed error between real and predicted peak KAM reduction. To assess if there was a reduction of accuracy when testing the model on new data, we compared the mean signed errors of the synthetic training data, synthetic test data, and Carnegie Mellon test data. After testing for normality using the Kolmogorov-Smirnov test, we compared the mean signed errors with a one-way analysis of variance (ANOVA) test and post-hoc Tukey’s Honestly Significant Difference (HSD) test for multiple comparisons. We used a significance level of 0.05 for all statistical tests.

## 3. Results

### 3.1 Learned gait patterns

Synthetic toe-in KAM correctly captured that all subjects reduced their first KAM peak. The group mean of the first KAM peak was 3.065 %BW*HT with 95% CI [2.15, 3.98] during baseline gait and 2.622 %BW*HT with 95% CI [1.71, 3.53] during toe-in gait. The mean predicted first peak of the KAM was 2.681 %BW*HT with 95% CI [1.76, 3.61] (Fig 4). Predicted knee joint center and center of pressure trajectories were more accurate in the mediolateral direction than in anterior-posterior; the resulting synthetic KAM trajectory was estimated with an RMSE of 0.253 %BW*HT (±0.112 %BW*HT) (Table 2).

**Table 2.**
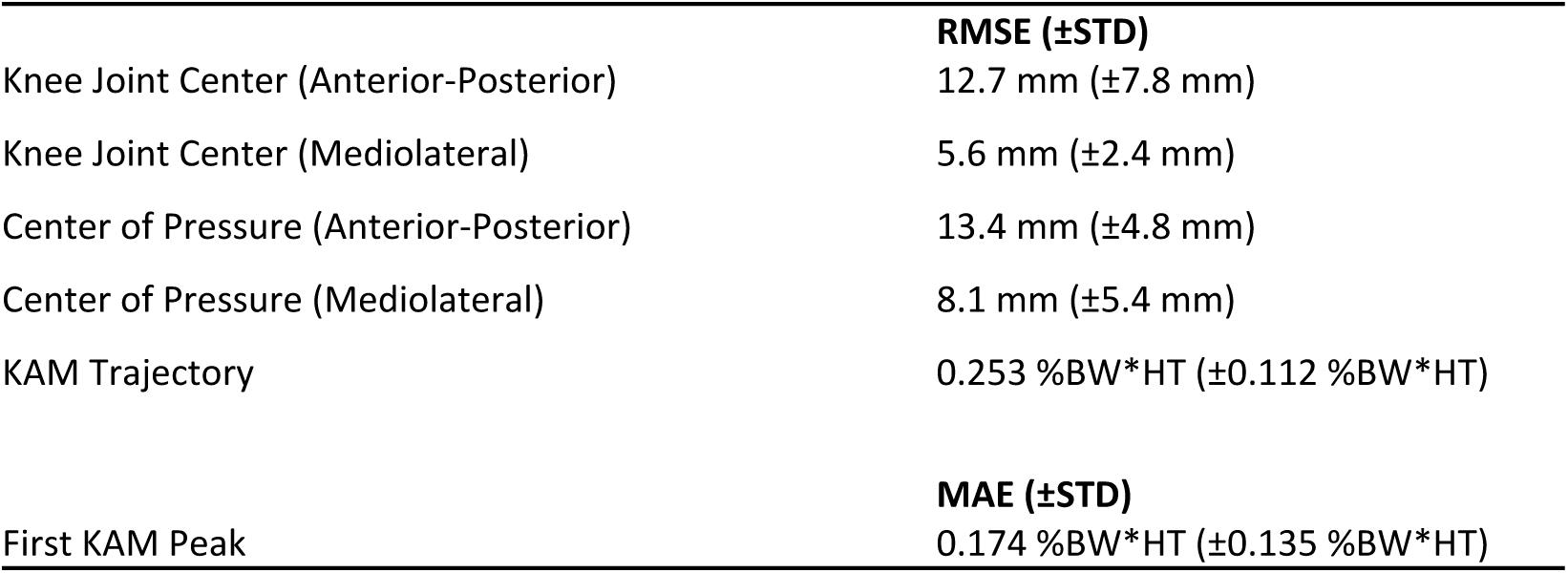
Evaluation of the learned gait patterns against ground truth measurements in the Stanford dataset

**Fig 4.**
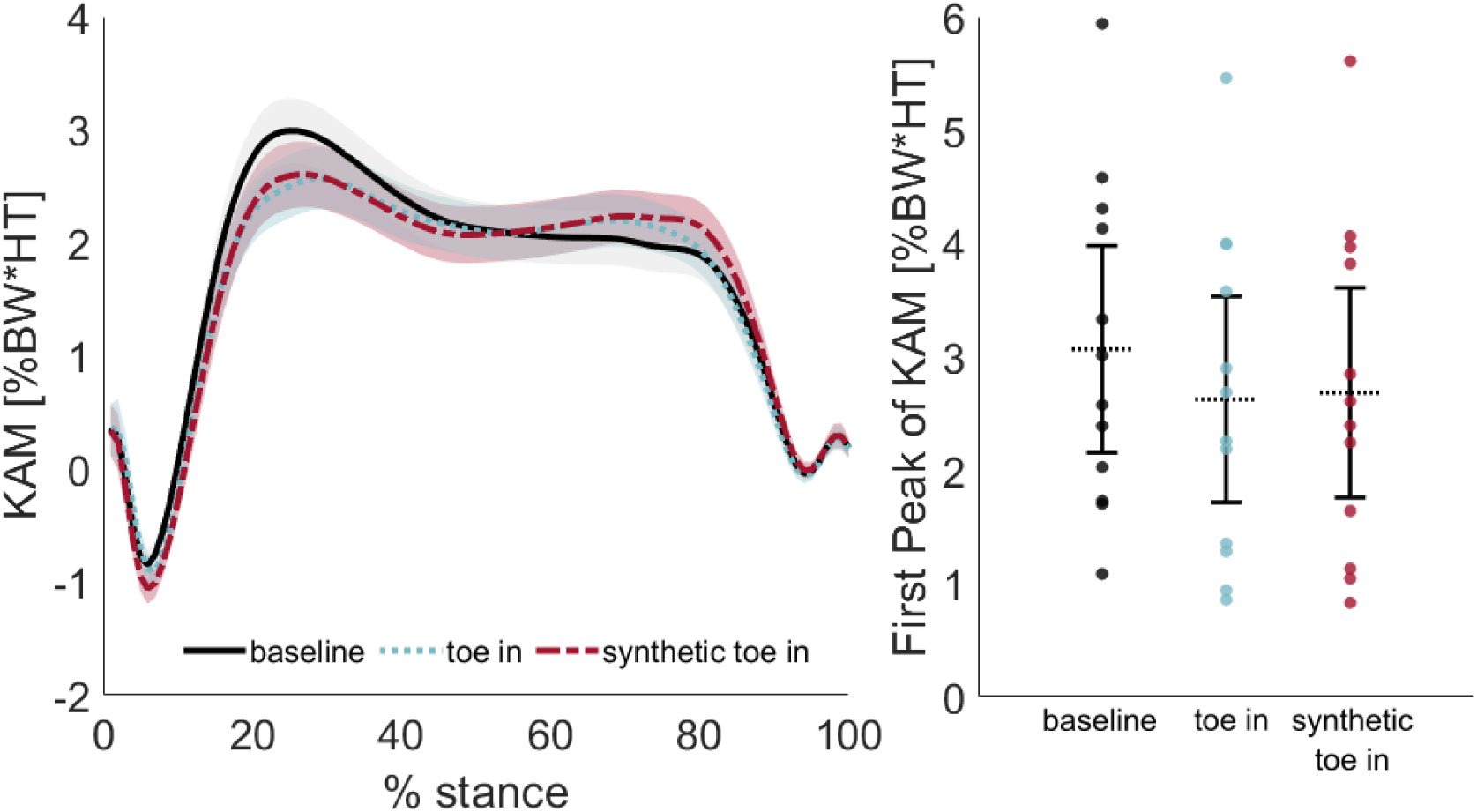
Mean KAM and first peak accuracy across subjects (Stanford dataset). With leave-one-out cross-validation, the synthetic toe-in KAM trajectory (red, dashed line) from the Stanford dataset closely matched the ground-truth toe-in KAM (blue, dotted line). Synthetic KAM captured the within-subject reduction in the first peak of KAM relative to baseline (black, solid line). The left plot captures mean (±STD) KAM trajectories across subjects, while the right plot shows individual and mean peak KAM with 95% confidence intervals.

### 3.2 Training and evaluating the predictive model

The predictive model estimated the first peak reduction with an MAE of 0.095 %BW*HT (±0.072 %BW*HT) (Fig 5). The average signed error across all toe-in angles for each of the 108 subjects in the training set was distributed around zero with 95% CI [-0.020, 0.020]. The mean signed error for the 15 subjects in the synthetic test set was also distributed around zero with 95% CI [-0.057, 0.061]. The model corresponding to the minimum MSE used all six features, with an alpha regularization term of 0.7. The toe-in angle was the strongest predictor, with a linear weighting coefficient of β1 = 0.30 (*p* < 0.0001). Increased valgus angle during static alignment (β2 = -0.015, *p* = 0.0002) and increased weight (β3 = -0.014, *p* = 0.0025) contributed to a smaller KAM reduction. Baseline foot progression angle (β4 = -0.0049, *p* = 0.198), height (β5 = -0.0041, *p* = 0.398), and walking speed (β6 = 0.0034, *p* = 0.375) did not significantly contribute to a reduction in KAM.

**Fig 5.**
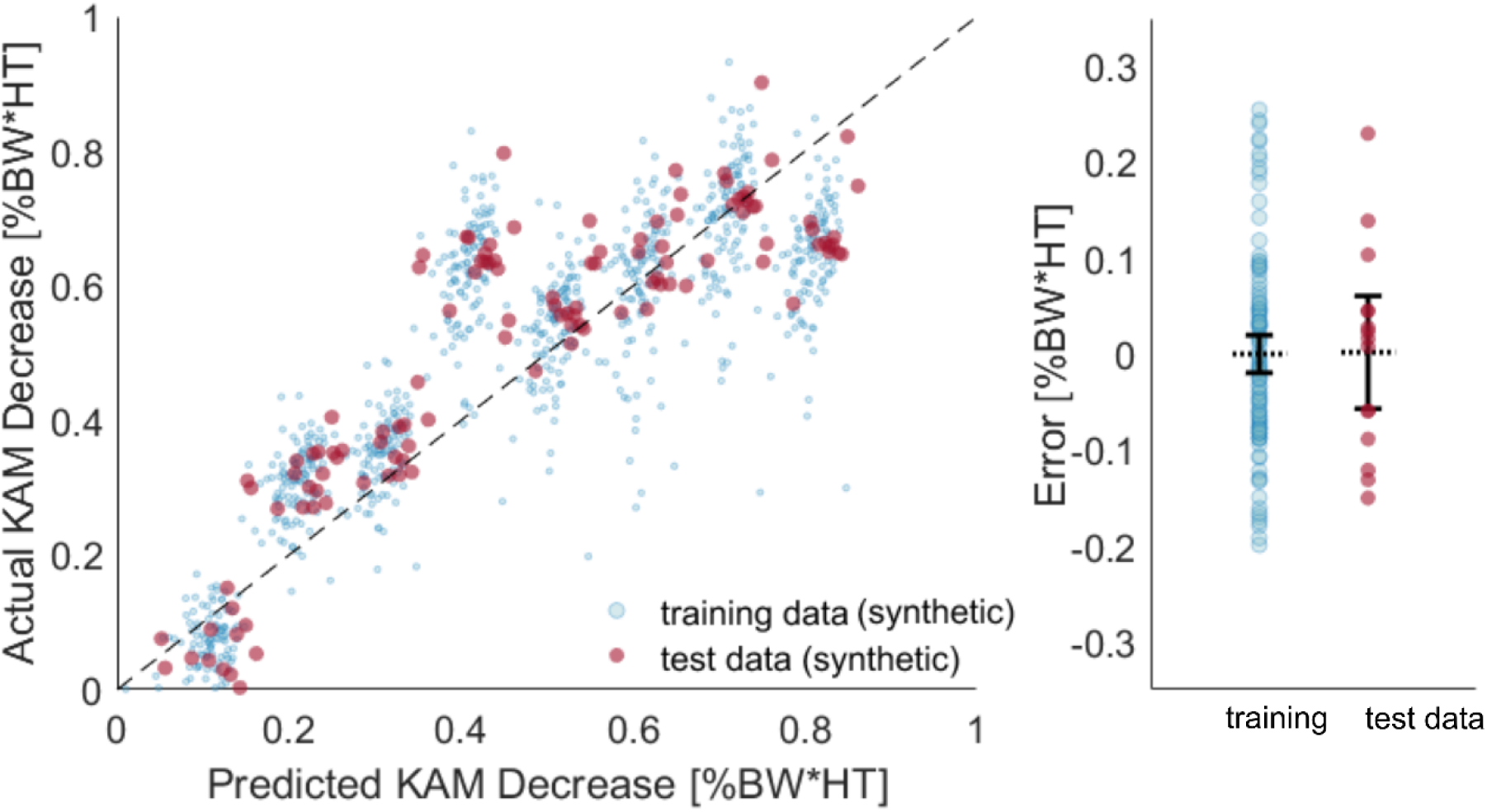
Prediction of KAM reduction using synthetic training data (Calgary Running Injury Clinic dataset). Synthetic toe-in data from 108 subjects were used to train the predictive model, which achieved an MAE of 0.0826 %BW*HT (± 0.0628 %BW*HT) on a synthetic test set of 15 subjects. Signed error (the difference between actual and predicted KAM reduction) of the training and test sets were similarly distributed around zero.

### 3.3 Validating the learned gait patterns and predictive model on a separate dataset

The synthetic toe-in KAM correctly captured that all subjects reduced the first KAM peak. The group mean of the first peak was 2.602 %BW*HT with 95% CI [2.08, 3.12] during baseline gait and 1.982 %BW*HT with 95% CI [1.47, 2.50] during toe-in. The mean predicted first peak of the KAM was 2.006 %BW*HT with 95% CI [1.46, 2.56] (Fig 6). The within-subject variation of the first peak of KAM was greater than the error in predicted KAM peak: across all steps for each subject, the average standard deviation of the first peak during baseline gait was 0.306 %BW*HT. The knee joint center and foot center of pressure toe-in gait patterns were predicted with similar accuracy to the Stanford dataset. The predicted knee joint center and center of pressure trajectories were again more accurate in the mediolateral direction than in anterior-posterior, and the resulting synthetic KAM trajectory was estimated with an RMSE of 0.335 %BW*HT (±0.121 %BW*HT) (Table 3).

**Table 3.** Validation of the learned gait patterns using the Carnegie Mellon dataset

| | RMSE (±STD) |
| --- | --- |
| Knee Joint Center (Anterior-Posterior) | 13.3 mm ($\pm 8.2$ mm) |
| Knee Joint Center (Mediolateral) | 4.5 mm ($\pm 2.7$ mm) |
| Center of Pressure (Anterior-Posterior) | 15.4 mm ($\pm 8.8$ mm) |
| Center of Pressure (Mediolateral) | 9.0 mm ($\pm 6.8$ mm) |
| KAM Trajectory | 0.335 %BW*HT ($\pm 0.121$ %BW*HT) |
| | MAE (±STD) |
| First KAM Peak | 0.170 %BW*HT ($\pm 0.101$ %BW*HT) |

**Fig 6.**
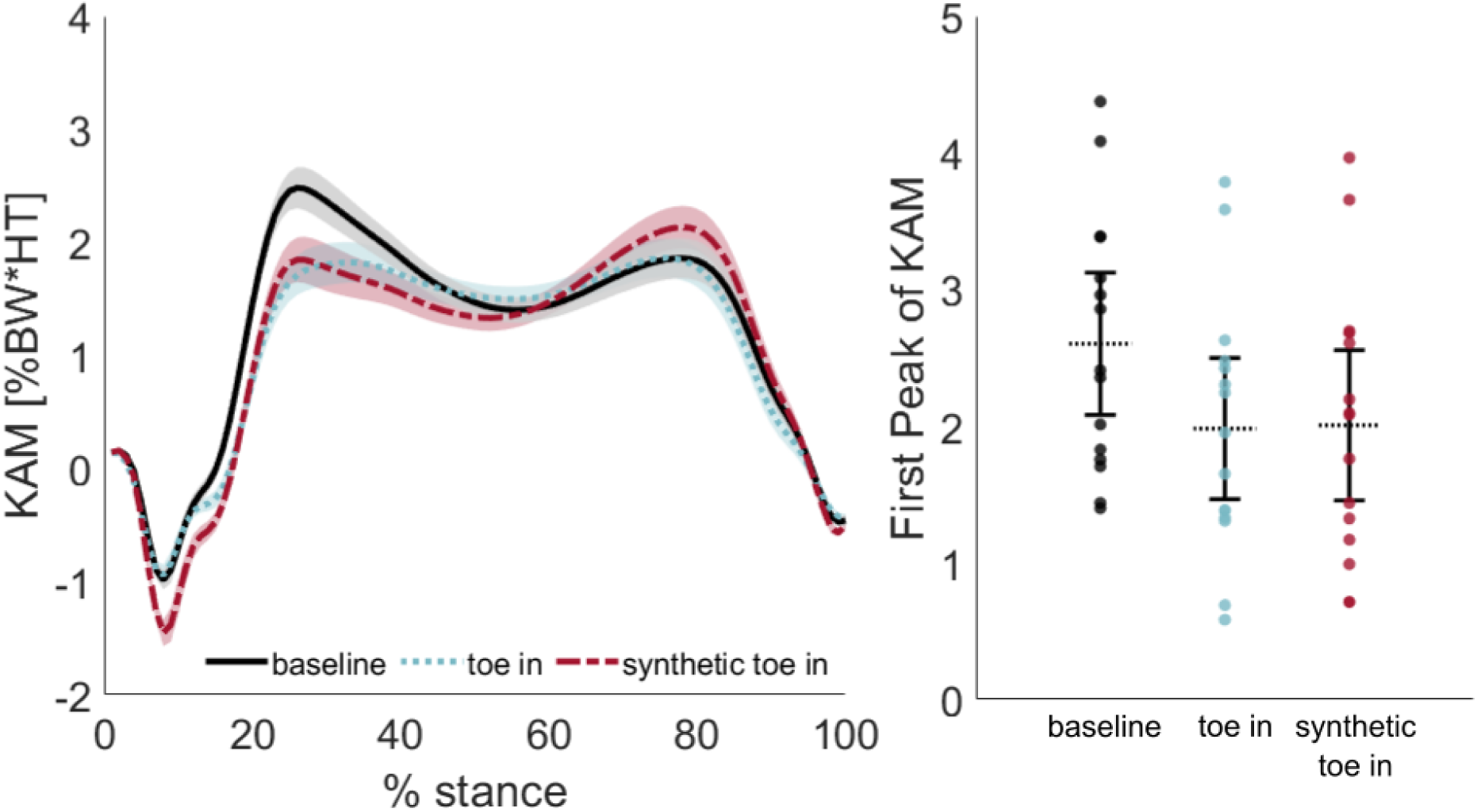
Validation of synthetic toe-in KAM (Carnegie Mellon dataset). The synthetic toe-in KAM trajectory (red, dashed line) closely matched the real toe-in KAM (blue, dotted line). Synthetic KAM captured the within-subject reduction in the first peak of KAM, relative to baseline (black, solid line). The left plot captures mean (±STD) KAM trajectories across subjects, while the right plot shows individual and mean peak KAM with 95% confidence intervals.

The predictive model trained on the synthetic data could estimate first peak KAM reduction of the Carnegie Mellon dataset with an MAE of 0.134 %BW*HT (±0.0932 %BW*HT) (Fig 7). The mean [95% CI] signed error between the predicted and real first peak KAM reduction was 0.068 %BW*HT [-0.017, 0.15]. Using only the toe-in angle, the strongest predictor, as a feature, KAM reduction was estimated with an MAE of 0.187 %BW*HT (±0.151 %BW*HT).

**Fig 7.**
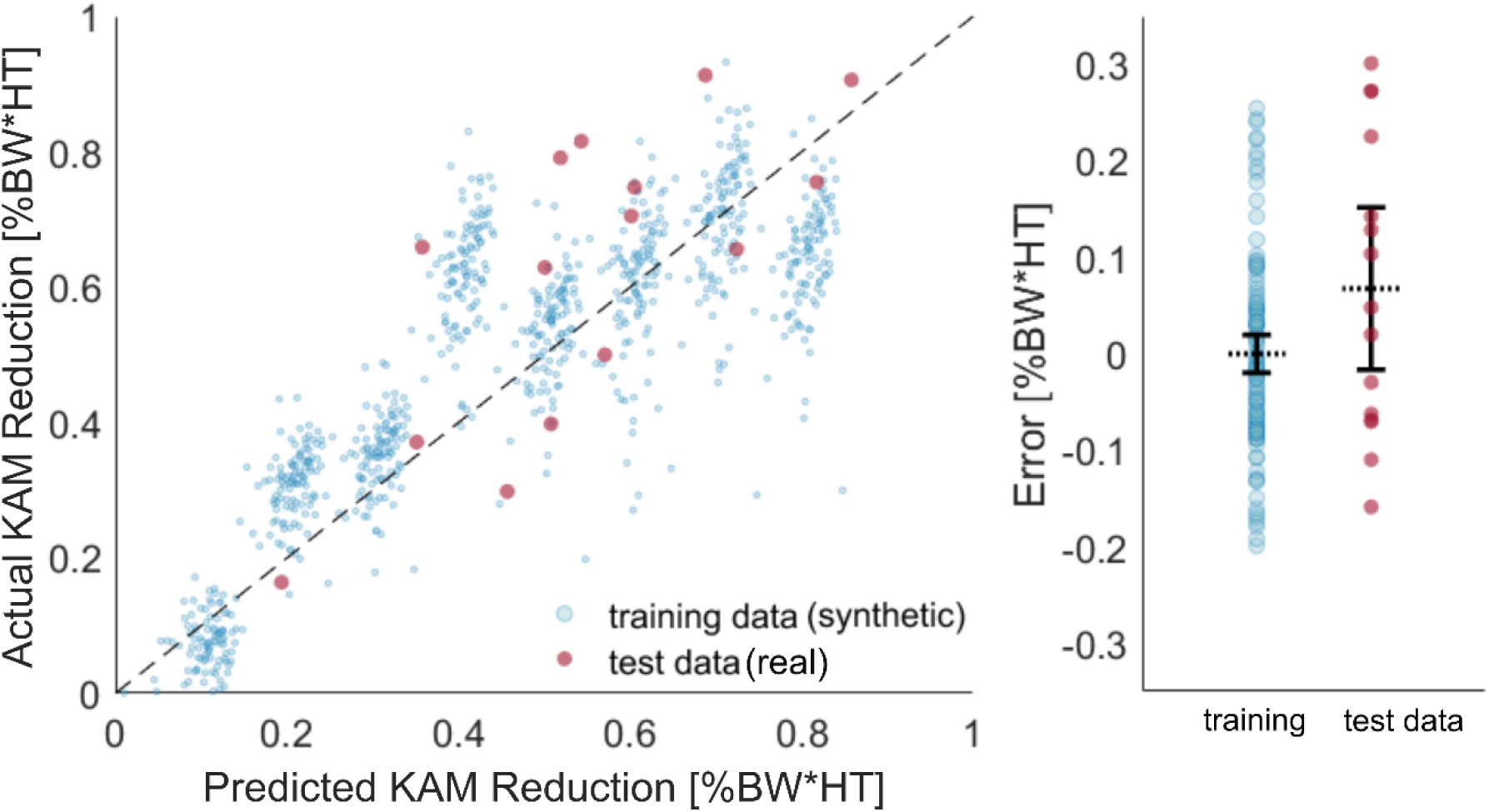
Validation of the predictive model (Carnegie Mellon dataset). An independent dataset of toe-in gait from 15 subjects was used to evaluate the predictive model, which achieved an MAE of 0.134 %BW*HT (±0.0932 %BW*HT). The mean signed error of the model was not significantly different between the Carnegie Mellon test data and the synthetic training or synthetic testing data.

Although the mean signed error of the independently collected data differed from the synthetic training and testing data, these differences were not statistically significant (*p* = 0.082). We found no statistically significant difference between the mean signed error of the synthetic training and synthetic test data (*p* = 0.998), synthetic training and real test data (*p* = 0.0635), or synthetic test data and real test data (*p* = 0.224).

## 4. Discussion

The aim of this work was to enable prediction of the first peak KAM reduction with a toe-in gait modification using features easily obtained in the clinic. Gait retraining is not yet standard in clinical practice, in part because a full gait analysis is required to identify a foot progression angle that reduces the KAM. With this predictive model and emerging lightweight haptic feedback systems [29], we hope that clinicians may one day be able to prescribe and initiate therapeutic gait retraining in a matter of minutes. As a practical evaluation of the model, we obtained a test error of 0.134 %BW*HT on data collected in a separate gait laboratory, falling within the range of the subjects’ step-to-step KAM variation (0.306 %BW*HT). These results may help to make toe-in gait modifications, based on gait retraining protocols, more widely accessible to clinicians.

Several study characteristics and limitations should be considered when interpreting the reported findings. In this study, KAM was computed using the lever-arm calculation method. Although the inverse dynamics link-segment method is sometimes preferred when real-time KAM estimates are not imperative [6], both methods show significant agreement in assessing percent change with a gait intervention, with a mean (±STD) difference of 5% (±14.1%) [30]. When synthesizing toe-in KAM, a 5% error in estimating toe-in peak KAM reduction would have resulted in a difference of 0.0298 %BW*HT, which is an order of magnitude less than the Carnegie Mellon subjects’ step-to-step KAM variation (0.306 %BW*HT). More importantly, using the lever arm method enabled us to synthesize toe-in KAM across datasets that used different marker sets. Additionally, some subjects in the Stanford dataset increased the second peak of the KAM with toe-in, resulting in a falsely predicted second peak increase for the Carnegie Mellon dataset. However, the average estimated second peak increase was smaller than the estimated first peak reduction, resulting in an overall smaller KAM impulse, which may still provide therapeutic benefit [31]. *Toe-out* gait modifications typically aim to reduce the second KAM peak [17], so future addition of toe-out gait data may facilitate more accurate prediction of first and second peak changes. Finally, as all of the subjects in the Carnegie Mellon dataset reduced their first KAM peak with toe-in, it is not clear if the model could accurately predict a clinical non-responder [32, 33]. While within the range of reported values [13, 17, 34], the Carnegie Mellon dataset average reduction in first peak KAM (0.62 ± 0.226 %BW*HT) trended toward being greater than that of the Stanford dataset (0.44 ± 0.242 %BW*HT). This difference may be due to disparities in pain-free mobility between healthy and osteoarthritic cohorts, as well as the method of toe-in gait retraining: here, we provided vibration feedback based on the toe-in angle, whereas the Stanford researchers provided feedback on the tibial angle in the frontal plane, indirectly inducing a toe-in gait that reduces KAM. The predictive model, built upon patterns learned from a cohort with a smaller KAM reduction, slightly underestimated the real KAM reduction (mean signed error = 0.068 %BW*HT), although we did not find this difference to be significant (*P*=0.082). Incorporation of several representative datasets from multiple laboratories would further the generalizability of such a model.

KAM reduction with toe-in gait can be predicted within the range of values seen in baseline gait by relying on a limited set of features. The toe-in angle positively correlated with KAM reduction, which is consistent with the previous finding that foot progression angle is a significant predictor of change in KAM [13]. However, using *only* toe-in angle as a predictor estimated KAM reduction to an MAE of 0.187 %BW*HT (± 0.151 %BW*HT), compared to 0.134 %BW*HT (±0.0932 %BW*HT) using the full set of 6 features. The next most salient predictor was that an increased valgus angle during static alignment was related to a smaller KAM reduction, supporting previous findings that KAM reduction is greater in participants with more varus alignment [34]. Although KAM was normalized to height and body weight, an increase in weight was predictive of a smaller KAM reduction. As knee osteoarthritis patients at a higher weight have a larger weight-normalized KAM than their lean age-matched osteoarthritic controls [35], further investigation is warranted to determine the effect, if any, of weight on KAM reduction and gait retraining outcomes. In a previous study, a neural network trained on 3D anatomical features from 86 subjects was used to classify whether an individual would increase or reduce their first peak KAM with toe-in or toe-out gait modifications, attaining accuracies of up to 85% [36]. The most salient features of their model included those related to the position of the pelvis, knee angle in the frontal plane, and sway of the trunk; in the future, a sensor-fusion approach that combines wearable sensing and advances in markerless motion capture could make use of these additional pelvis and trunk features to improve our predictive model. However, while our model was capable of correctly predicting a KAM reduction for all subjects in the external dataset, we instead sought to estimate the *extent* of KAM reduction, rather than *classify* by whether KAM would decrease or not. Providing an estimate of the KAM reduction with gait retraining may empower clinicians to weigh the expected benefit of retraining against that of other treatment alternatives.

The accuracy of the knee joint center and center of pressure predictions were within the error range of joint center location estimates obtained with optical motion capture. Although divergent experimental protocols prevent meaningful meta-analysis [37], estimates of the knee joint center position have been found to vary from 14 mm to up to 40 mm [38-40]. As compared to gold-standard position tracking with intra-cortical bone pins and MRI, optical motion capture suffers from skin movement artifacts and inconsistency in marker placement across subjects, experimenters, and laboratories [41]. The inter- and intra-dataset validation of the learned gait patterns showed comparable accuracy between the Stanford and Carnegie Mellon datasets, giving us further confidence in this method of generating the synthetic KAM data. The use of these validated toe-in gait patterns may therefore enable reliable predictions of changes in joint kinetics by any research group, without the need for additional motion capture trials.

Enabling accurate prediction of an expected gait change without collecting baseline kinetic data is a significant step toward bringing insights from the gait lab into the clinic. Collecting gait data with altered foot progression angles is a time-consuming iterative process: after becoming acquainted with the biofeedback paradigm, subjects must acclimate to each new gait pattern, at which point their kinetics may be evaluated. In the absence of universal guidelines on short- and long-term gait retraining learning rates [42], experimenters allow subjects a minimum of 2 minutes to acclimate to the new foot progression angle, with 10-30 minutes of training to enforce the modified gait strategy [15-18]. With the use of the predictive model, this experimental time may be reduced drastically: height, weight, and static knee alignment are already commonly measured in the clinic, whereas wearable sensors can one day allow walking speed and baseline foot progression angle to be standard vitals that comprise a patient’s unique gait health profile [43].

A growing acceptance of modern data science methods is transforming how gait biomechanics is analyzed by providing researchers with opportunities to gain insight from highly variable and complex measurements [44]. These methods require large quantities of heterogeneous data to avoid overfitting and to generalize well to unseen subjects [45]. Although biomechanists have begun to advocate for the need to share data and for the importance of normative databases [46,47], these datasets do not yet exist for more specialized needs such as gait retraining. Here, we demonstrate one promising method to overcome the present paucity of data by generating a synthetic dataset based on limited ground-truth examples. With continued exploration of synthetic data generation, we hope to move toward a future in which the therapeutic benefit of a potential treatment can be assessed to determine a custom treatment path for any patient.

In conclusion, this work demonstrates that it is possible to predict the extent to which a patient will benefit from gait retraining therapy using clinically available measures. It also illustrates the feasibility of training predictive models with synthetic data. By further harnessing the growing capabilities of emerging portable biomechanics labs that fuse sensing and vision approaches, these models may empower clinicians to prescribe gait retraining therapy in the clinic.

## Acknowledgments

The authors would like to thank Owen Pearl, as well as the Stanford University and Calgary Running Injury Clinic researchers for their assistance with data collection. They would also like to thank the participants for their contribution to this study.

